# Projected warming disrupts embryonic development and hatch timing in Antarctic fish

**DOI:** 10.1101/2025.05.28.656708

**Authors:** Margaret Streeter, Nathalie R. Le François, Thomas Desvignes, Jacob Grondin, John H. Postlethwait, H. William Detrich, Jacob M. Daane

## Abstract

Rising ocean temperatures pose significant threats to marine ectotherms. Sensitivity to temperature change varies across life stages, with embryos often being less tolerant to thermal perturbation than adults. Antarctic notothenioid fishes evolved to occupy a narrow, cold thermal regime (−2 to +2°C) as the high-latitude Southern Ocean (SO) cooled to its present icy temperatures, and they are particularly vulnerable to small temperature changes, which makes them ideal sentinel species for assessing climate change impacts. Here, we detail how predicted warming of the SO may affect embryonic development in the Antarctic bullhead notothen, *Notothenia coriiceps*. Experimental embryos were incubated at +4°C, a temperature projected for the high-latitude SO within the next 100–200 years under high emission climate models, whereas control embryos were incubated at present-day ambient temperature, ∼0°C. Elevated temperature caused a high incidence of embryonic morphological abnormalities, including body axis kinking/curvature and reduced body size. Experimental embryos also developed more rapidly, such that they hatched 68 days earlier than controls (87 vs. 155 days post-fertilization). Accelerated development disrupted the evolved timing of seasonal hatching, shifting larval emergence into the polar winter when food availability is scarce. Transcriptomic analyses revealed molecular signatures of hypoxia and disrupted protein-folding in near-hatching embryos, indicative of severe cellular stress. Predictive modeling suggested that temperature-induced developmental disruptions would narrow seasonal reproductive windows, thereby threatening population viability under future climate scenarios. Together, our findings underscore the vulnerability of Antarctic fish embryos to higher water temperature and highlight the urgent need to understand the consequences of disruption of this important trophic component on ecosystem stability in the SO.

**Significance Statement:** Antarctic fishes evolved cold-adapted phenotypes suited to the stable thermal conditions of the Southern Ocean, yet are threatened by rising temperatures. The impact of rising temperatures on early life stages in Antarctic fishes is not well understood; our findings show that projected warming may induce premature hatching, developmental abnormalities, and molecular stress responses in embryos, potentially reducing recruitment and leading to population instability and trophic-level ecosystem disruptions. These results underscore the urgency of assessing climate-driven vulnerabilities across life stages of Antarctic marine organisms to refine population projections and enhance conservation strategies amid ongoing environmental change.

## Introduction

Although the Southern Ocean (SO) has historically been one of the most thermally stable marine habitats, it is projected to experience dramatic environmental changes (1). Between 2005 and 2017, the SO absorbed 45-62% of global ocean heat despite covering only about 25% of the ocean surface (1). Sea surface temperatures along the West Antarctic Peninsula have risen by 1°C since 1955 (2), with projections suggesting an additional warming of 3-4°C within the next 100-200 years under high-emission scenarios (SSP5-8.5; **Fig. 1A**)(3). Moreover, Antarctic sea ice cover has rapidly declined since its recent peak in 2014, losing as much ice in three years as the Arctic did over three decades, indicating a shift toward a new and warmer climate regime (4, 5). Understanding the impact of these changes on Antarctic marine organisms is crucial for forecasting future ecosystem dynamics.

**Fig. 1.**
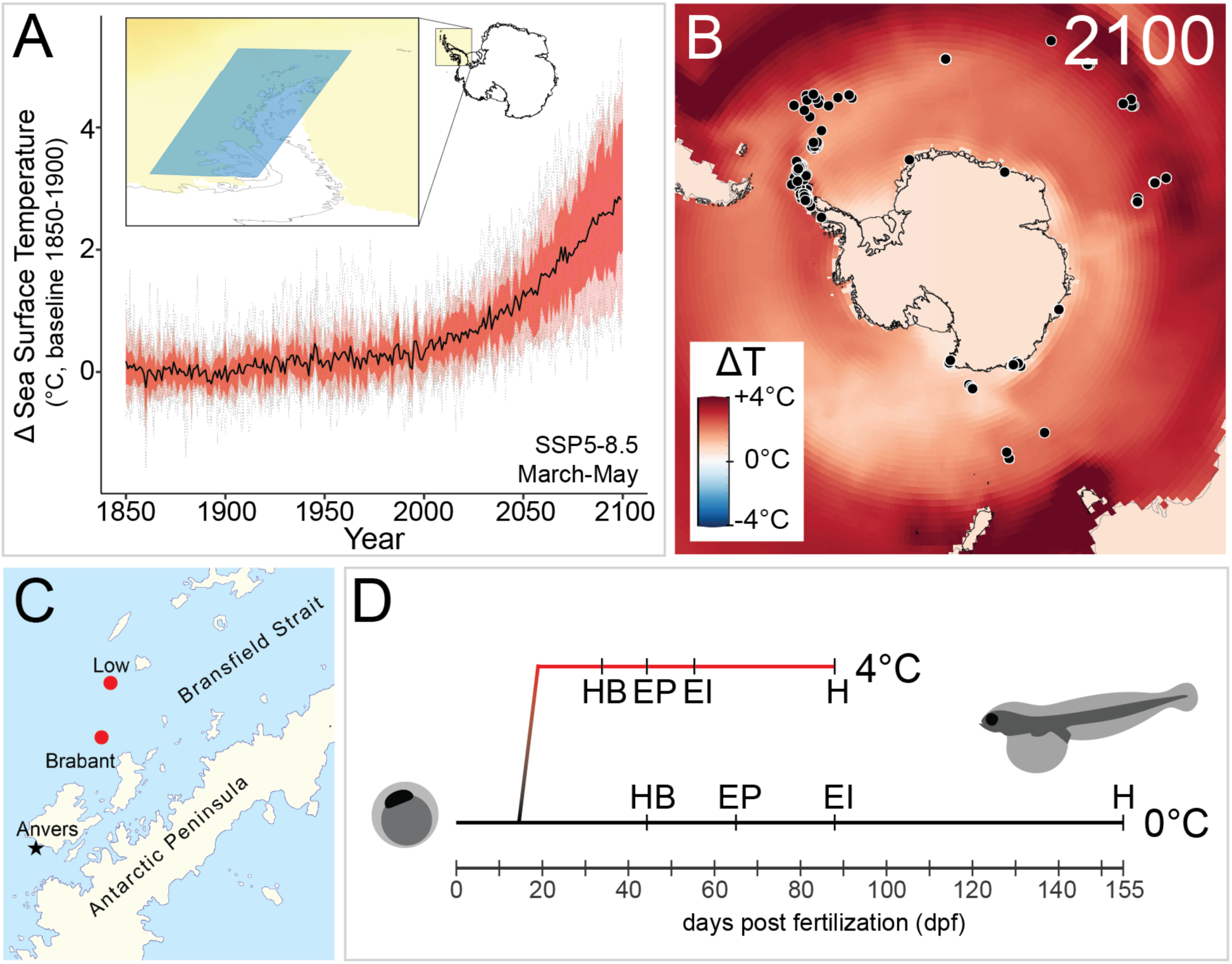
Southern Ocean (SO) warming and *Notothenia coriiceps* embryonic development. (A) Projected change in sea surface temperature (SST) along the Antarctic peninsula (inset) by 2100 and during *N. coriiceps* breeding season (March-May) under the SSP5-8.5 scenario, based on the CMIP6 dataset accessed via the Copernicus Interactive Climate Atlas (3). The black line indicates median SST projection, while the gray lines indicate individual climate models. Red and pink shading reflect 25-75th and 10-90th percentiles, respectively. (B) Projected SST changes in the SO under SSP5-8.5 in 2100, overlaid with historical *N. coriiceps* catch records (black dots) from AquaMaps (118). Temperature heatmap from the IPCC Interactive Climate Atlas (119). (C) Fishing locations where *N. coriiceps* specimens used in this study were collected (red dots) and Palmer Station where embryos were raised using flow through seawater from neighboring Arthur Harbor (black star). (D) Developmental timeline of embryos incubated under ambient (0°C) and elevated (4°C) temperatures, highlighting stages selected for RNA sequencing: HB (heartbeat), EP (50% eye pigmentation), EI (eye iridescence), and H (first hatching).

The ichthyofauna of the modern SO is dominated by species of the Notothenioidei suborder (order Perciformes) (6). Before the mid-Miocene, Antarctic fish biodiversity significantly decreased along with the cooling and glaciation of Antarctica, leading to local extinction of most fish taxa (6). With competition reduced, the benthic common ancestor of notothenioids radiated adaptively about 10 million years ago to yield over 100 species that exploit all niches in the SO (6–9). Today, notothenioids constitute about 90% of fish biomass on the High Antarctic continental shelf, 66.5% of species captured in the Scotia Sea, and include keystone species that are essential for maintaining ecosystem function (6, 10–13).

Persistently cold SO temperatures (−2 to +2°C annually (14)) have contributed to specialized biochemical, cellular, and physiological traits in notothenioids, including antifreeze glycoproteins, loss of red blood cells in icefishes, and absence of a typical heat shock response (8, 15, 16). These and other cold specializations, however, contribute to narrow thermal tolerances in adults and limit their capacity to cope with thermal stress (17–19). While much research has focused on the responses of adult notothenioids to future warming (20–24), the effects of elevated temperatures on other life stages, especially embryos, remain poorly understood.

Thermal resilience varies throughout a fish’s life cycle, with spawning adults and embryos often exhibiting the lowest tolerance ranges (17). Temperature has pleiotropic effects on embryos by influencing the kinetics of biochemical reactions, metabolic rates, protein folding and stability, oxidative stress, and sex determination (25). Because the responses of different cell types to thermal stress vary, developmental asynchronies and teratological effects may occur (26). Furthermore, thermal stress can redirect limited yolk resources away from growth and development toward fueling cellular stress responses, resulting in stunted growth and reduced larval fitness (27–29).

Depending on the climate scenario, an estimated 10-60% of all fish species are projected to exceed their embryonic developmental temperature limits within their current ranges by 2100 (17). However, experimental data on thermal tolerance limits are scarce for most fish embryos, making it difficult to accurately model the effects of future climate change across diverse fish lineages. The buoyant embryos of several notothenioid species may be particularly vulnerable to thermal stress because, in the absence of sea ice, they are exposed to environmental fluctuations near the sea surface (30). Previous studies of Antarctic embryo thermal resilience are sparse and limited to short-term thermal exposures of field-collected embryos from a single Antarctic dragonfish species (*Gymnodraco acuticeps*, Bathydraonidae) (31, 32). Temperature effects on developmental viability varied from limited to strong, potentially due to the timing of heating or other variables (31, 32).

In the present study, we examine the development of the Antarctic bullhead notothen, *Notothenia coriiceps*, in the context of projected SO warming over the next 100-200 years. We show that increased temperature during embryonic development shortens the time to hatching, causes morphological abnormalities, impacts the phenology of hatching, and perturbs gene expression related to hypoxia, protein homeostasis, and the cellular stress response. Using these insights, we predict reduction in seasonal embryonic survival and larval recruitment with potential shifts in the timing of breeding.

## Results

### Development of N. coriiceps embryos under rising ocean temperatures

For environmental conditions, we chose 4°C, predicted to be reached by 2100-2200 given the unmitigated climate emission scenario Shared Socioeconomic Pathways (SSP5-8.5)(1, 3)(1, 3, 33). (**Fig. 1A**). We focused on the Antarctic bullhead notothen, *Notothenia coriiceps*, an abundant species (**Fig. 1B**, (34)) with a known breeding season and an established developmental staging series (35, 36). Although *N. coriiceps* adults are benthic (36), their zygotes are buoyant, form part of the zooplankton, and are exposed to changing regional sea surface temperatures; thus *N. coriiceps* serves as an excellent proxy for multiple notothenioid species with pelagic embryos (35, 37).

Eighty adult *N. coriiceps* were collected during their Austral fall breeding season from two locations along the Antarctic peninsula (**Fig. 1C**; see *Materials & Methods*). After transport to the aquarium facilities at Palmer Station, Antarctica, fish were maintained in flow-through seawater aquaria at ∼0°C. Males and gravid females were injected with Ovaprim®, a gonadotropin-releasing hormone analog, to stimulate ovulation and spermiation. Four spontaneous spawning events over three days (5/25-5/27/2018) in two 2.5-m^3^ circular tanks (30 fish/tank) produced approximately 44,000 embryos. Embryos from these four spawns were pooled, divided into two equivalent technical replicate groups, and incubated at the ambient water temperature of Arthur Harbor (controls, ∼0°C). On day 15 post fertilization, corresponding to ∼40% epiboly, each replicate group was split to give one cohort to be incubated at +4°C (experimental) and one (control) at ambient temperature (**Fig. 1D**, **S1, S2**). To avoid abrupt heat shock to the embryos, we applied a temperature ramp of ∼1°C/day to the two experimental replicates over days 15-19 to increase long term survival chances (**Fig. S3** shows the temperature profiles measured for experimental and control incubators). The delayed ramp was intended to mitigate the role of maternal effects on our results. We estimate that the maternal-to-zygotic transition in *N. coriiceps* embryos occurred between 5-9 dpf based on cell number, the change to asynchronous and uneven cell division, and the onset of epiboly in early gastrulation (35), as found in other fish species (38). Thus, our findings reflect the impact of warming on transcription of the zygotic genome.

**Fig. 2.**
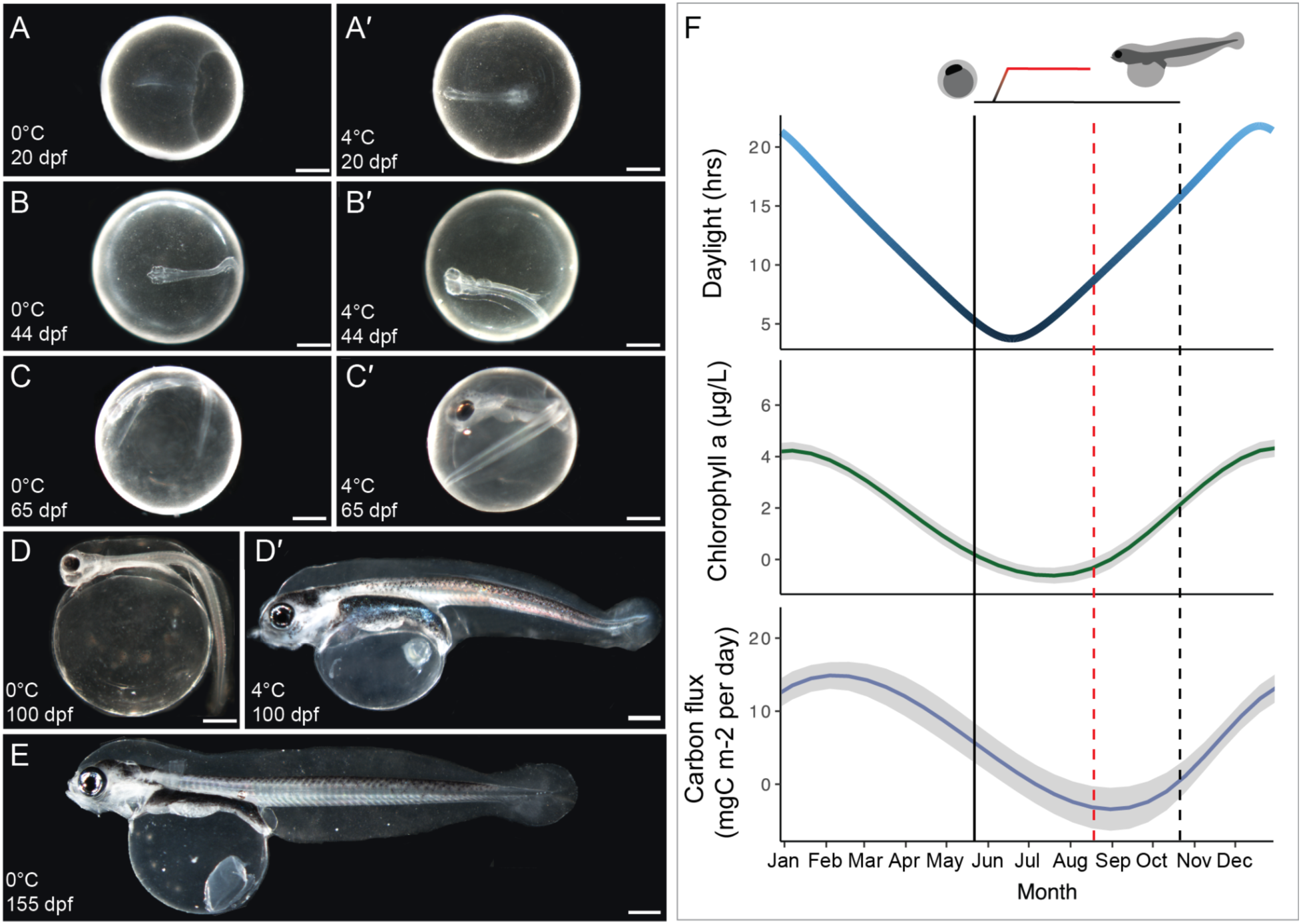
Accelerated developmental rate and phenological asynchrony in *N. coriiceps* embryos raised at elevated temperature. Developmental progression of *N. coriiceps* embryos from 20 days post fertilization (dpf) to hatching at 0°C (A-E) or at 4°C (A’-D’). Scale bars represent 1 mm. (F) Comparison of relative hatch timing in *N. coriiceps* at 0°C (black dashed line) and 4°C (red dashed line), overlaid with environmental data from the Palmer Station Long-Term Ecological Research (LTER) database on Anvers Island for chlorophyll *a* and carbon flux (39–41). The chlorophyll *a* dataset is condensed from 30 years of measurements, and the carbon flux data is comprised of 20 years of sediment trap data (gray areas represent confidence bands of localized regression (loess, geom_smooth()).

The survival rates of embryos were similar between treatments, with the exception of an early mortality event (37.9% mortality over the first 40 days) in one of the ambient technical replicates (**Fig. S4A,B**) that may have resulted from a decline in water quality due to accumulation of dead embryos. Subsequently, embryonic survival at 4°C at 74 dpf was 93.2% and 92.8% for the two replicates, compared to 96.3% and 93.1% for the 0°C replicates at 118 dpf (**Fig. S4B**). These results indicated that mortality across treatments was unlikely to affect the results reported below.

### Accelerated development of N. coriiceps embryos led to hatching during austral polar winter

During the 15-day incubation at 0°C, embryos in the four cohorts progressed through cleavage, epiboly, and establishment of the embryonic axis on the *N. coriiceps* staging series (29) (data not shown). Following the temperature ramp up, control and experimental embryos completed gastrulation, segmentation, organogenesis, and entered skeletogenesis prior to hatching, but those at 4°C attained these milestones earlier (**Fig. 1D, 2A-E, S5**). Segmentation in experimental embryos began at 20 dpf at 4°C versus 23 dpf at 0°C, first heartbeat occurred at 34 dpf at 4°C compared to 44 days for controls, and the onset of retinal pigmentation and iridescence for the experimentals appeared at 44 and 65 dpf, versus 55 and 87 dpf for controls (**Fig. S5**; **Table S1**). Thus, development of *N. coriiceps* embryos was substantially accelerated at 4°C compared to ambient temperature, but embryos in both treatments nonetheless passed through the same developmental stages.

Control embryos had just begun hatching at 155 dpf but had not yet reached a peak prior to the conclusion of the field season, which coincided with the onset of hatching (**Fig. 2E**). In contrast, embryos raised at +4°C began hatching at 87 dpf, and hatching continued through 100 dpf (**Fig. 2D’**). Thus, hatching of experimental embryos was completed at least 55 days earlier compared to first hatching of controls. This phenological asynchrony has important implications for post-hatching larval survival and development: hatching at 155 dpf coincides with the arrival of polar spring (November), when food availability increases along the Antarctic Peninsula (**Fig. 2F**; see carbon flux west of Anvers Island and chlorophyll *a* for Arthur Harbor at Palmer Station (39–41)), whereas experimental larvae would hatch in late winter (August-September), a period of low primary productivity and comparatively limited food availability (42).

### Development at 4°C resulted in morphological abnormalities in larval stages

Embryos raised at 4°C exhibited several gross morphological abnormalities. Approximately 65% of randomly-sampled, near-hatching embryos showed abnormalities at 4°C, compared to only 20% at 0°C (**Fig. 3A-F**). The abnormalities included increased body-axis curvature and deformation (**Fig. 3C**, which are phenotypes previously observed in fish embryos under thermal stress (26)), and craniofacial abnormalities involving the jaw (**Fig. 3D,E**). The most common abnormalities involved embryos with kinked, bent, and shortened tails.

**Fig. 3.**
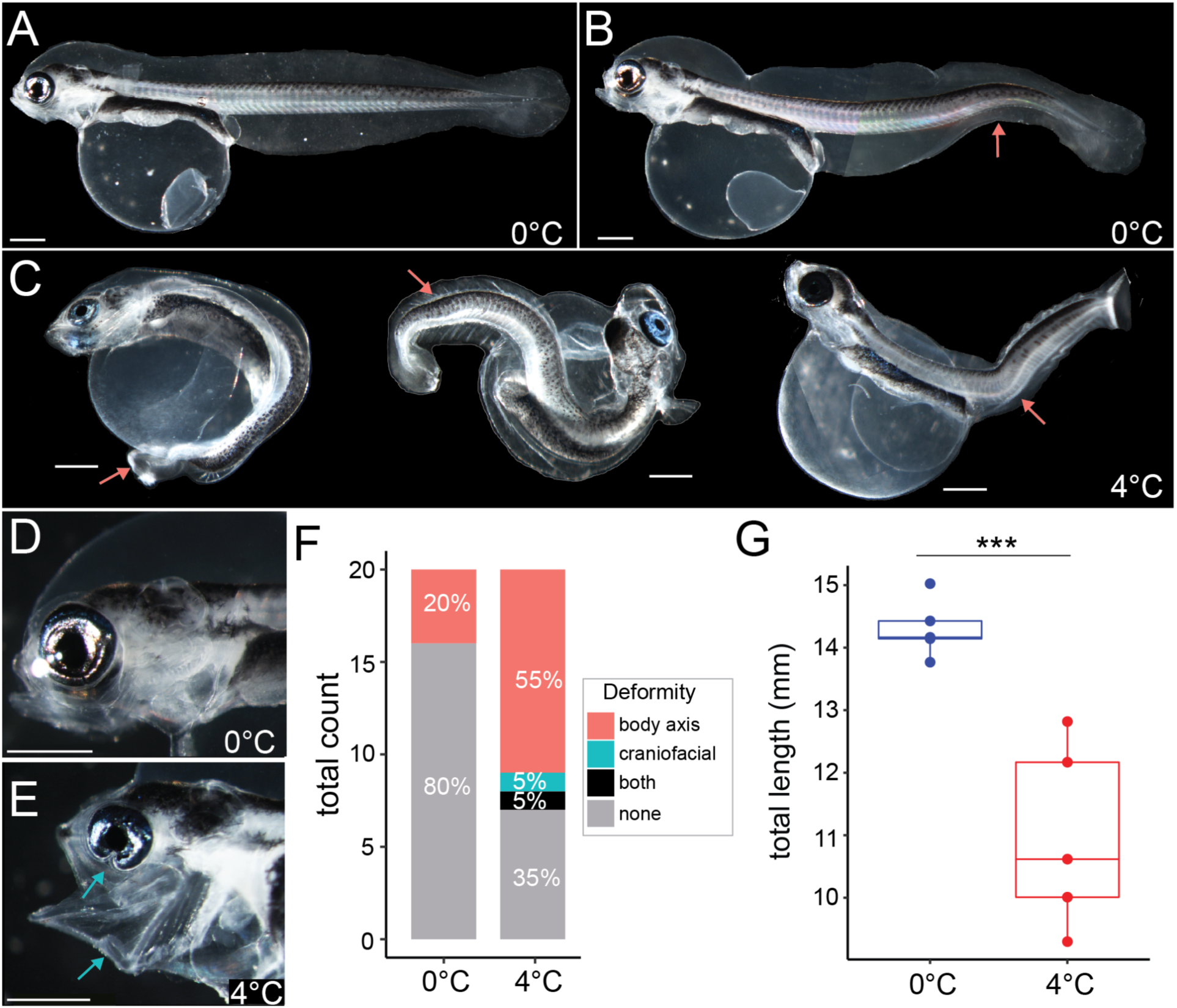
Elevated rates of morphological abnormalities in *N. coriiceps* embryos developing at 4°C. (A) A recently hatched larva developed at 0°C showing normal morphology. (B) An example of an abnormal larva raised at 0°C with body axis curvature. Arrow pointing to curvature. (C) Examples of abnormalities at hatching-stage embryos raised at 4°C. Scale bars in (A-C) represent 1 mm and arrows point to specific abnormalities including a failure of the axis to fully elongate and axis curvature. (D-E) Comparison of craniofacial morphology between a normal larva reared at 0°C (D) and an abnormal larva reared at 4°C (E). Scale bars represent 1 mm and arrows point to a defect in the ventral part of the eye and the fixed open mouth. (F) Quantification of deformities in near-hatch embryos reared at 0°C and at 4C. (G) Total length just before first hatching at 0°C (149 dpf) and at 4°C (86 dpf) (**Table S2**). *** indicates significance with a Wilcoxon rank sum exact test, p-value = 0.008.

Near hatching, experimental embryos were shorter than control embryos, with 4°C animals measuring 10.98 mm ± 1.47 at 86 dpf (mean ± SD) compared to ambient water controls measuring 14.30 mm ± 0.47 (149 dpf) (**Fig. 3H**, see also **Fig 2.D’**, **E**; **Table S2**). All hatchlings still had large yolk sacs that remained after the yolk was used, which has been observed in *N. coriiceps* (35, 42) (**Fig. 2D’**, **E**; **Table S2**).

### Development at elevated temperature altered the transcriptomic profiles of embryos and hatchlings

To evaluate the effect of elevated temperature on *N. coriiceps* development at the molecular level, we performed bulk RNA sequencing on individual embryos collected at the morphological milestones previously described: heartbeat onset (HB), 50% eye pigmentation (EP), eye iridescence (EI), and hatching (H). We aligned sequencing reads to the genome assembly of close relative *Notothenia rossii* (GenBank ID: GCA_949606895.1, (43)) rather than to the *N. coriiceps* genome assembly (GCA_000735185.1, (44)) because of the greater contiguity (contig N50: 383.4 kb vs. 17.5 kb) and BUSCO completeness (94.9% vs. 77.3% single copy orthologs) of *N. rossi*’s genome assembly. Differential gene expression (DGE) was analyzed using both DESeq2 and EdgeR, which apply geometric normalization and trimmed mean of M values normalization, respectively (45, 46). In total, we estimated DGE for 21,172 genes that passed gene-level filtering criteria out of 24,432 annotated genes. Genes were considered significantly differentially expressed only if they were significant by both methods at FDR-adjusted p ≤ 0.05. **Fig. S6** shows that DESeq2 (47) clustered sample expression profiles by developmental stage rather than temperature treatment, which indicates that comparable transcriptional profiles were recovered by our morphology-based sampling approach.

We found that hundreds of genes were highly differentially expressed (log2FoldChange > 1 or < −1) at each developmental stage: 1226 at heartbeat onset (HB), 940 at 50% eye pigmentation (EP), 651 at eye iridescence (EI), and 859 at first hatching (H), **Fig. 4A**, **Table S3**). The variability of gene expression between the two temperature treatments was similar for most of development, but at hatching, more genes showed a qualitative increase in non-Poisson noise at 4°C compared to 0°C (**Fig. 4B**). To our surprise, most differentially expressed genes in at least one of the studied developmental stages (2,231 of 2,881) were specific to a single developmental stage (**Fig. 4C**). Only 34 genes were differentially expressed at all stages (FDR-adjusted p ≤ 0.05), eight of which had high expression differences (log2FoldChange ≥ 1 or ≤ −1) (**Table S3**). Six of these eight were *apolipoprotein B a* (*apoba*), *caveolin 4 a* (*cav4a*), *4-hydroxyphenylpyruvate dioxygenase* (*hpdb*), *insulin-like growth factor binding protein acid labile subunit* (*igfals*), *keratin 4* (*krt4*), *switching B cell complex subunit 70 a* (*swap70a*). The remaining two genes lacked prior annotation but were identified by BLAST as homologous to *b cell lymphoma 2 like 16* (*bcl2l16)* and *formin-like protein 20* (*LOC104963861, N. coriiceps*) (**Table S3**).

**Fig. 4.**
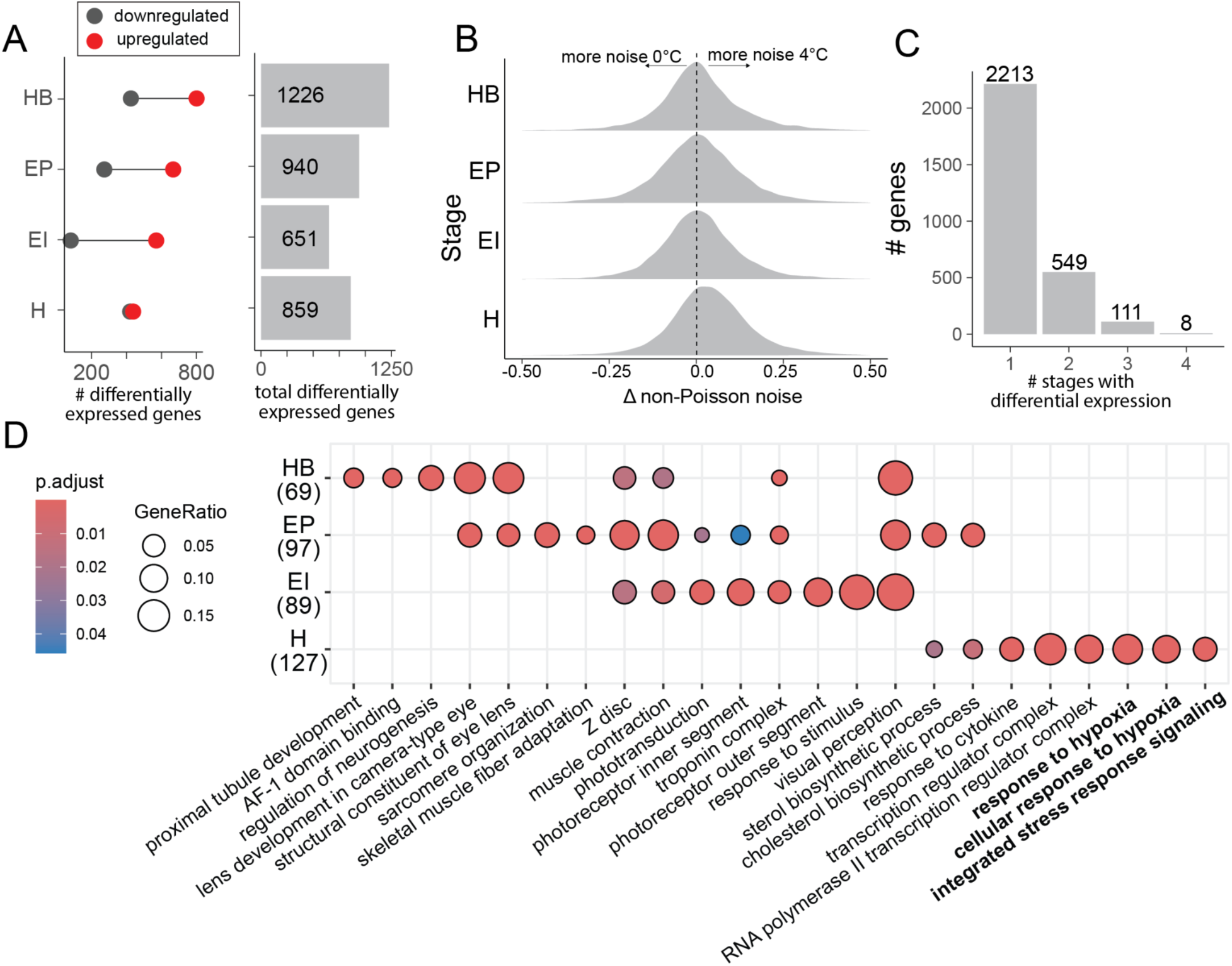
Patterns of differential gene expression during *N. coriiceps* development. (A) Number of genes significantly differentially expressed at each stage (padj ≤ 0.05, LFC ≤ −1 or ≥ 1). Red dots indicate genes upregulated at 4°C, gray dots indicate genes downregulated at 4°C relative to 0°C, and gray bars represent the total number of differentially expressed genes at each stage. (B) Distribution of the difference in non-Poisson noise for each gene between treatments (GAMLSS)(110). Positive values indicate greater transcriptional variance at 4°C versus 0°C. (C) Number of genes differentially expressed at one, two, three, or all four developmental stages. (D) Dot plot of Gene Ontology (GO) enrichment over developmental time. The top six enriched GO terms at each stage are shown. Dot size represents the proportion of genes within a GO term that are differentially expressed, while color indicates statistical significance (FDR-adjusted p). For (A-D), developmental stages include first heartbeat (HB), 50% eye pigmentation (EP), eye iridescence (EI) and first hatching (H). Numbers in parentheses indicate the total number of significantly enriched terms of the upregulated genes at each stage.

We then investigated whether differentially expressed genes were enriched for specific biological functions using the enricher function in ClusterProfiler v4.14.4 (48). Genes upregulated at 4°C showed significant enrichment for 69 gene ontology (GO) terms at heartbeat (HB), 97 at 50% eye pigmentation (EP), 89 at eye iridescence (EI), and 127 at first hatching (H) stages (FDR-adjusted p ≤ 0.05, **Table S4**). The GO terms enriched for upregulated genes at HB were related to vision, neurogenesis, and muscle development; at EP to sterol biosynthesis and muscle function; at EI to visual perception, stimulus response, and troponin complex; at H to stress response, hypoxia, and transcription (**Fig. 4D**, **Table S4**). Genes downregulated at 4°C were enriched for 16 GO terms at HB, 78 at EP, 10 at EI, and 119 at H (**Table S4**). The GO terms for downregulated genes at HB were linked to transmembrane transport; at EP to Notch signaling and neurogenesis; at EI to interneuron migration; at H to glucuronidation, steroid metabolism, and hemostasis (**Fig. 4D**, **Table S4**). Most enriched GO terms were specific to one stage, none occurred across all stages, and hypoxia-related terms appeared only at hatching (**Fig. 4D**, **Table S4**).

We anticipated that temperature elevated beyond evolved thermal limits would disrupt developmental signaling due to the pleiotropic disruption of most cellular processes (25), and due to the presence of morphological abnormalities (**Fig. 3**). However, we found that only the Notch signaling pathway was consistently enriched (both up- and downregulated) in the GO enrichment analysis and was dysregulated at all stages except EI. Other pathways (Bmp, Fgf, Wnt, Hedgehog) were not enriched for expression dysregulation, although a few individual developmental pathway members were differentially expressed (**Table S4**).

### Transcriptomes of thermally stressed larvae exhibited molecular signatures of hypoxia and cellular stress

We hypothesized that stress-related genes would be consistently upregulated at +4°C throughout development. Results, however, showed increased transcriptional noise and enrichment for cellular stress responses (e.g., hypoxia, oxidative stress, and mitochondrial dysfunction) in heated embryos compared to controls only at hatching (**Fig. 4B**, **D**; **Table S4**). Several highly upregulated genes at hatching are involved in the cellular response to hypoxia, including the key oxygen sensors *egl-9 family hypoxia inducible factor 2* (*egln2*, also known as *phd1;* see *Materials and Methods*) and *egl-9 family hypoxia inducible factor 3* (*egln3*/*phd3*), which act upstream of *hypoxia-inducible factor 1-alpha* (*hif1a*) signaling (**Fig. 5B,C**; **Table S4**) (49). Downstream Hif targets involved in anaerobic metabolism (e.g., *ldha*, *pfkfb3/pfk2*) were also upregulated (**Fig. 5A,B,C**; **Table S4**). Upregulation of these glycolytic genes would be expected to enhance anaerobic metabolism under low oxygen conditions (50).

**Fig. 5.**
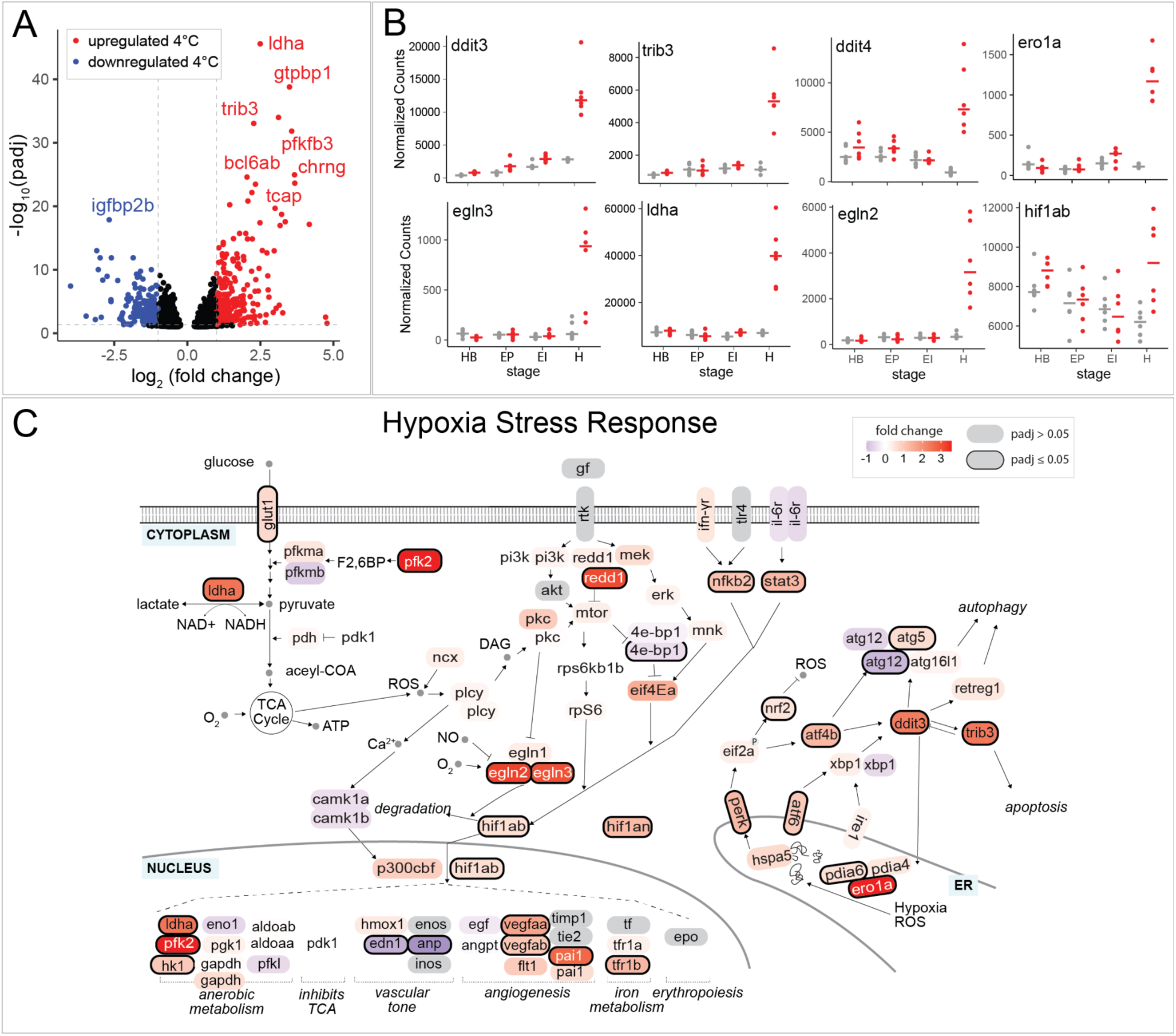
Differential expression of hypoxia and integrated stress response genes during the hatching of *N. coriiceps* embryos. (A) Volcano plot showing differentially expressed genes at hatching. Blue dots represent genes with greater expression at 0°C (padj ≤ 0.05, LFC ≤ 1), while red dots denote genes with greater expression at 4°C (padj ≤ 0.05, LFC ≥ 1). (B) Expression profiles of selected genes throughout development. Developmental stages include heartbeat (HB), 50% eye pigmentation (EP), eye iridescence (EI) and first hatching (H). Normalized counts are from DESeq2. (C) Diagram of the hypoxia stress response pathway based on KEGG (NT06542, NT06534) (120–122). Each oval is a gene in this pathway, with color indicating log₂ fold change. Genes significantly differentially expressed (padj ≤ 0.05) are outlined in bold. Genes not identified in the dataset are shown in gray and genes with multiple annotated copies in the *N. rossii* genome are represented twice.

The unfolded protein response (UPR) is activated by hypoxia, which disrupts oxidative protein folding in the endoplasmic reticulum (ER) (51). Several core UPR genes were upregulated at hatching at +4°C: *reticulum oxidoreductase 1 alpha* (*ero1a*) and *protein disulfide isomerase family A member 6* (*pdia6*) encode proteins involved in oxidative protein folding.

*Eukaryotic translation initiation factor 2 alpha kinase 3* (*eif2ak3*/*perk*) and *activating transcription factor 6* (*atf6*) encode proteins that serve as initial sensors of protein folding stress. *DNA damage-inducible transcript 3* (*ddit3*/*chop*) encodes a transcription factor whose downstream targets include *ero1a* and *tribbles pseudokinase 3* (*trib3*) (**Fig. 5B**,**C**; **Table S4**). *ero1a* and *trib3*, which are involved in autophagy and apoptosis in response to protein folding stress (52–54), were two of the most upregulated genes at hatching at +4°C. Finally, *DNA damage inducible transcript 4* (*ddit4*/*redd1*), a negative regulator of translation that functions downstream of various cellular stress response pathways (55), was also upregulated during hatching at +4°C (**Fig 5B,C**; **Table S4**). Together, the enrichment data suggest substantial hypoxic and proteostatic cellular stress in heated embryos at hatching but not before hatching.

## Discussion

### Developmental teratologies in heated embryos

Modest warming of the Southern Ocean is likely to disrupt embryonic development of its stenothermal fauna, whether vertebrates or invertebrates. Our findings with *N. coriiceps* align with thermal tolerance studies of Antarctic invertebrates. Antarctic krill (*Euphausia superba*) embryos show reduced hatching success at temperatures of 3°C, and 50% of nauplii are abnormal at 5°C (56). Percent of normal developing Antarctic sea urchin *Sterechinus neumayeri* drops significantly at 48 hours post fertilization (hpf) with a small increase in temperature (80% normal developing at 1 °C vs. 30% normal developing at 3°C) (59). Similarly, Antarctic starfish (*Odontaster spp.*) embryos reach 20% non-viability at 3°C (57). Thus, slight oceanic warming is very likely to have profound impacts on embryonic development in SO species, whether vertebrate or invertebrate, with few species showing resilience.

We did not observe increased embryonic lethality at +4°C (**Fig. S4**), perhaps because we applied the temperature ramp after gastrulation, thus after highly temperature-sensitive developmental stages (58–62). Furthermore, we did not address other environmental stressors, such as increasing ocean acidification from rising pCO₂ or freshening of surface waters from melting ice and increased precipitation (1). Elevated pCO₂ reduces overall thermal tolerance in fish adults and embryos (31, 63–65), including the Antarctic dragonfish, *G. acuticeps* (31). Thus, mortality and malformation of *N. coriiceps* embryos under the IPCC scenario SSP5-8.5 involving multiple stressors would likely be more severe than we report here with a single stressor.

We observed a high incidence of body-axis curvature in the +4°C embryo population at hatching (**Fig. 3**), a common defect in fish embryos raised at supraphysiological temperature that has been linked to misfolded protein accumulation in notochordal sheath cells (26). The observed upregulation of UPR genes (e.g., *ddit3*, *atf6*, *perk*, and *ero1a*) in *N. coriiceps* +4°C hatchlings supports proteostatic stress as an important determinant underlying body axis curvature (**Fig. 5**), as seen in zebrafish (26). Increased transcriptional noise in hatchlings at +4°C (**Fig. 4B**) may reflect the diverse teratologies observed in this stage of embryogenesis (**Fig. 3C**) or may be driven by cell-type-specific responses to acute cellular stress near hatching (26). Body-axis defects, combined with reduced body size, likely decrease fitness in temperature-stressed embryos which would impact recruitment success (66).

After hatching, embryos transition from relying on yolk reserves to actively feeding, a critical period for survival in fish (67). Thus, body size at hatching significantly influences foraging success and starvation resistance (66). At +4°C, *N. coriiceps* embryos exhibited reduced growth compared to ∼0°C embryos (**Fig. 3**), consistent with a prior study of Atlantic cod subjected to thermal and pH stress (63). Decreased embryonic growth under stress is thought to reflect the prioritization of essential homeostatic functions over developmental growth (68).

One stressor, hypoxia, can induce premature hatching in fishes, leading to smaller larvae (60, 69–71). Our gene expression data in *N. coriiceps* suggested hypoxia-driven precocious hatching may have occurred at elevated temperature (**Fig. 5**).

### Timing of hatching and phenological implications

The timing of larval hatching is crucial for survival of zooplankton and for trophic dynamics in planktonic assemblages (72, 73). Temporal displacements between trophic levels pose challenges to larvae that feed on seasonal prey, thereby impacting fish distributions and food web dynamics (74, 75). Fish embryos possessing large yolk reserves typically have long incubation periods and may be able to delay hatching to align with favorable ecological conditions (76–79), though specific hatch-inducing triggers, like hypoxia, may disrupt timing relative to other signals such as light (80).

Notothenioid embryonic incubation periods range from one month in cool-temperate, Sub-Antarctic species to ten months in high-latitude Antarctic species (30). The developmental time to hatching (∼five months (**Figs. 1, 2**)) that we observed for *N. coriiceps* embryos raised at ambient Palmer Station (Arthur Harbor intake) temperature is consistent with previous reports of six months at Palmer Station (35), seven months at King George Island (81), and five months at Signy Island (82). *N. coriiceps* embryos hatch with nearly empty yolk sacs (35, 42) (**Fig. 2D’**,**E**), which indicates that they must feed soon after hatching to survive (30). As the larval abundance of Antarctic fishes are correlated with primary productivity (11, 83), phenological coupling of hatching with the onset of polar summer is likely crucial for their survival.

### Shifting reproductive windows in an Antarctic fish

Because temperature strongly influences developmental rates (**Fig. 2**), projected SO warming may uncouple the evolved synchronization between the timing of breeding seasons and the eventual hatching of larvae at a time of food availability. In our study, *N. coriiceps* embryos began hatching in 155 days at 0°C and in 87 days at +4°C. Limited by two temperature data points for *N. coriiceps* embryos, we used a linear model to project developmental rates under future climate scenarios along the West Antarctic Peninsula (**Fig. 6**). Note, data in cods suggest a negative exponential relationship between temperature and developmental rate, with a near linear increase in rate between −1°C and 4°C that asymptotically approaches what is likely a maximum developmental rate beyond 4°C (84). We assumed that temperatures ≥ 4°C result in non-viable embryos due to morphological defects or hypoxia and that larvae hatching before or during the polar winter are non-viable due to food scarcity. Under present-day conditions (ERA5 (14)), eggs are fertilized in May, embryos develop over winter, and larvae hatch in mid-October/November during the austral spring phytoplankton bloom (**Fig. 6A**).

**Fig. 6.**
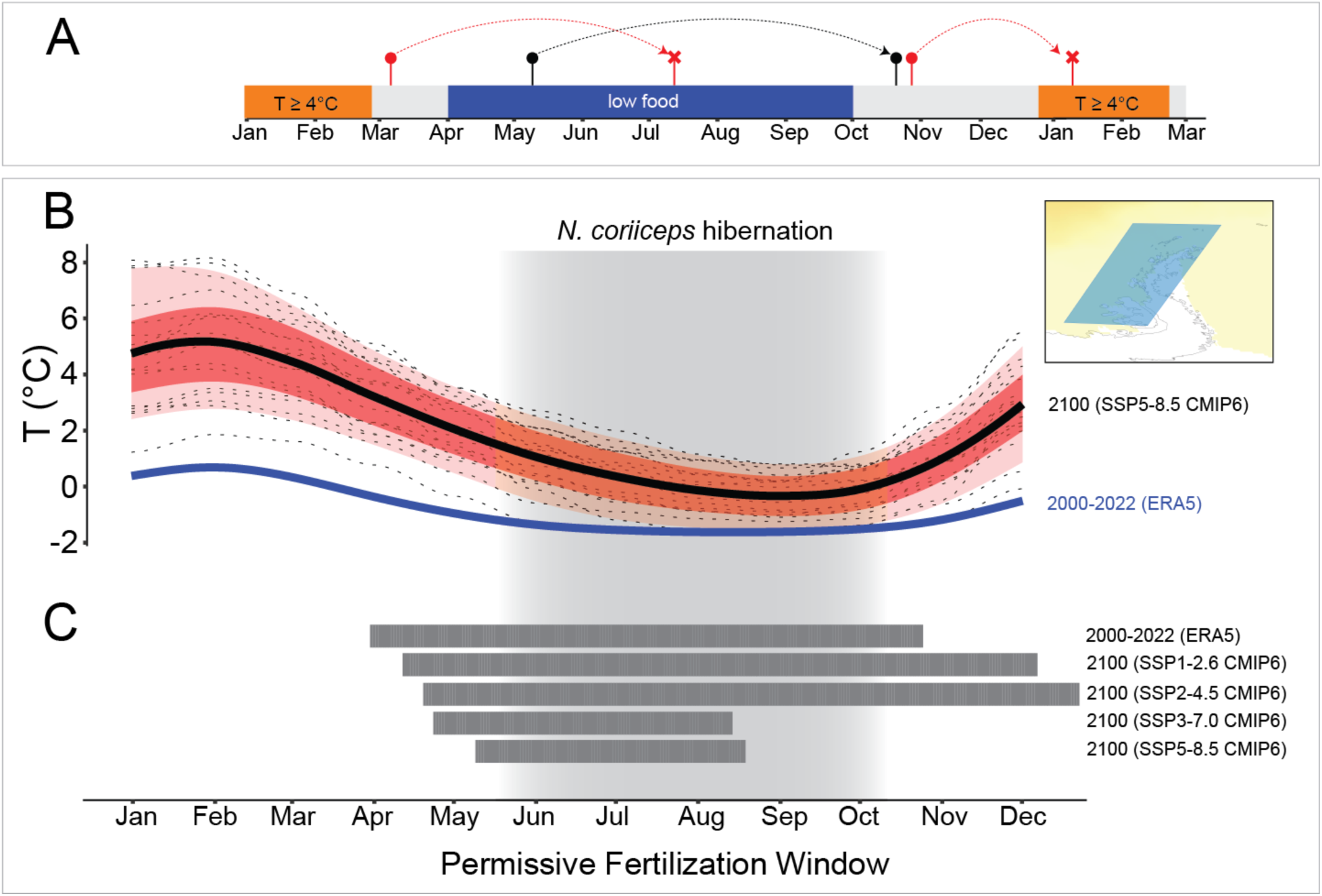
Modeling the timing of fertilization for *N. coriiceps* embryos in a changing climate. (A) Overview of development and hatch timing for *N. coriiceps* embryos. Fertilization is timed so that hatchlings emerge in polar spring. However, complications during development may arise if: 1) temperatures exceed 4°C, or 2) hatching occurs before or during the polar winter (April-October), when food availability is low. Red lines indicate examples of non-viable developmental windows from fertilization to hatching. Black lines indicate an example viable developmental window (B) Plot of monthly SST projections under a high carbon emissions scenario (SSP5-8.5), compared to the average monthly SST from 2000-2022 (ERA5). Dashed lines represent different climate models, the dark black line represents the smoothened median, and the red and pink shading indicates the 25-75th and 10-90th percentiles, respectively. (C) Bars represent fertilization dates that meet the criteria outlined in panel A. Approximate dates of hibernation for adult *N. coriiceps* are shaded in gray in panels B and C (85).

Under all future climate projections ((33), IPCC 2022), the window for successful fertilization shifts compared to current conditions (**Fig. 6B,C**). In lower emission scenarios, breeding season is lengthened by faster development, resulting in more opportunities for embryos that are fertilized in the spring to hatch outside of the winter period. However, in high emission scenarios, the temperatures rise too much, resulting in developmental problems. By 2100, winter temperatures should still support *N. coriiceps* embryonic development (**Fig. 6B,C**), such that late breeding into winter could improve embryonic survival by delaying hatching until spring. However, *N. coriiceps* adults exhibit hibernation-like behavior during winter, including reduced heart rate, metabolism, movement, and growth (85), which could make winter breeding improbable unless hibernation is temperature sensitive. Shifting breeding to spring (November) would lead to embryonic hatching during lethal austral summer conditions that exceed 4°C (**Fig. 6A**, red curve on right). *N.coriiceps* ability to adapt to future climate change may be determined by whether there is sufficient genetic variation or phenotypic plasticity in breeding behaviors to overcome hibernation-like activity or the if individuals have the ability to regulate or delay the timing of hatching to synchronize with phytoplankton bloom schedules.

### Hypoxia during thermal stress

Our transcriptomic findings support the hypothesis that hypoxia is a key factor limiting *N. coriiceps* embryo thermal tolerance (**Figs. 4**, **5**). Warmer temperatures accelerate biochemical reactions, resulting in an increased oxygen demand that may not be met by the fixed rate of oxygen diffusion across the chorion (61, 86–90). Larger eggs, common in cold climates, further increase anoxia risk since metabolic rate scales with egg size (30, 88, 91). As temperatures rise, polar fish embryos are thus likely at an elevated risk for hypoxia.

Embryo thermal tolerance shifts throughout development. In teleosts, oxygen consumption peaks at gastrulation and at hatching, but is generally lower during mid-development, aligning with periods of highest temperature-induced mortality (59–62, 92, 93). For example, the dragonfish *G. acuticeps* in −1 to −0.5°C water at McMurdo Sound can briefly tolerate temperatures above 8°C after gastrulation but show mortality at just 2°C if heating commences during gastrulation (31, 32). Consistent with these observations, we observed molecular signatures of hypoxia only at hatching and not at earlier developmental stages where oxygen demand is less (**Figs. 4,5**).

### Adaptability of Antarctic fishes to warming

The temperature changes modeled in this study are predicted to occur gradually over multiple generations. In other fish clades, populations and closely related species can differ significantly in embryo thermal tolerance (e.g., (84, 94)), indicating that thermal tolerance can be an evolvable trait. Notothenioids are characterized by slow generation times often exceeding 5-15 years, with *N. coriiceps* reaching maturity in about 5-7 years (36, 95, 96). These slow generation times would be predicted to limit their capacity to rapidly adapt to changing climates (97, 98).

However, several notothenioid lineages originally from Antarctic waters have successfully adapted to warmer conditions north of the polar front (6, 99). One example is the Maori chief (*Paranotothenia angustata*), a congener of *N. coriiceps*, which inhabits warmer coastal waters around New Zealand and Australia. Compared to *N. coriiceps*, *P. angustata* has secondarily evolved higher thermal tolerance in adults (100). Little is currently known about *P. angustata* reproduction and embryogenesis. Studying how these temperate-adapted notothenioids cope with warming as embryos would help project responses of Antarctic fishes to climate change and identify mechanisms driving the evolution of thermal tolerance.

### Summary

Our data revealed that projected temperature increases [4°C, by 2100-2200, unmitigated climate emission scenario Shared Socioeconomic Pathways (SSP5-8.5) (1, 3)] over the next 100-200 years are likely to severely impact *N. coriiceps* embryo development, causing high rates of morphological abnormalities, molecular stress responses, and asynchrony between developmental rates and seasonal environmental conditions. Although a +4°C sea surface temperature increase is currently on the extreme end of climate projections, transient seasonal variability or marine heatwaves from a higher baseline temperature could disrupt reproduction in *N. coriiceps* and other Antarctic fish. Similar studies across the life cycle of other organisms with diverse egg and embryo characteristics may provide key data points needed to improve accuracy of predictive climate change models.

## Materials and Methods

### Fish collection, maintenance, and spawning

Adult *N. coriiceps* specimens were collected south of Low Island and west of Brabant Island (Dallman Bay) along the West Antarctic Peninsula between April 20 and May 28 of 2018 (**Fig. 1C**). Eighty specimens were captured by deploying Otter trawls and baited traps from the *ARSV Laurence M. Gould* as previously described (101). Fish were maintained onboard ship in six 1 m^3^ flow-through isothermic tanks (Xactics, Cornwall, Ontario, Canada) with supplemental aeration. The fish were transferred within two days to the Palmer Station aquatic laboratories, where they were held in 2.5-m^3^ circular tanks supplied with flow-through, filtered, and aerated seawater from Arthur Harbor, as previously described (102). Sexually mature adults were housed at an average 25 fish per tank.

To promote gonadal maturation in captivity, males and females received up to two intraperitoneal injections of salmon gonadotropin releasing hormone (GnRH) analog as an ovulating and spermiating agent at a dosage of 0.5 mL/kg (Ovaprim Syndel, Ferndale, WA, USA). Tanks were checked several times a day for signs of spontaneous broadcast spawning, such as floating eggs or large amounts of foam on the surface due to tank aeration interacting with protein in the water. In total, there were 14 spawning events over 12 days. The first four spawning events were pooled to create the group of embryos used in this experiment, with 0 days post-fertilization (dpf) designated as the date most eggs were fertilized (two spawns on 5/26/2018 versus one each on 5/25/2018 and 5/27/2018). Spawning most likely involved mixed parentage as multiple males and females could have released gametes. All procedures were performed accordance with the Animal Care and Use Committee (IACUC) at Northeastern University (#15-0207R).

### Embryo culture

Embryos were cultured in two flow-through vertical incubation systems operated at Palmer Station (Marisource, WA, USA) in a walk-in refrigerated room kept at 2°C (**Fig. S1**). Seawater pumped directly from Arthur Harbor was first sand-filtered, then sterilized with UV light, and dispatched into two 50-L LLDPE reservoir tanks (Nalgene, Thermo Fisher Scientific, USA).

Water in the reservoir tanks was oxygenated with air pumps, before being distributed into the top section of the incubation systems (**Fig. S1**). One of the two reservoir tanks was heated to 4 ± 0.2°C using three feedback-controlled submersible heaters, while the other one was kept at ambient temperature. Oxygen levels were measured at the bottom of both incubator towers using a handheld multiparameter meter (Pro2030™ Dissolved Oxygen/Conductivity Meter (YSI Inc. Yellowspring, OH, USA)) and remained relatively stable (102.08 ± 0.72 % saturation at 0°C and 100.88 ± 0.53 saturation at 4°C) across incubator drawers (**Table S5**).

Fertilized *N. coriiceps* eggs averaged 4.36 ± 0.03 mm in diameter and 0.05 ± 0.002 g in wet weight. A total of 2.19 kg (∼44,000 embryos) were used, with 482 g placed in two incubator drawers at 0°C. At 15 days post-fertilization (dpf), 241 g of embryos per tray were transferred to an aerated incubator separate from the main incubation trays, then gradually heated by 1°C a day until reaching +4°C four days later at 19 dpf. Heated embryos were then moved to the heated vertical incubator system. Temperatures in both incubator systems were taken once an hour and averaged for each day using immersed temperature loggers inserted in the incubator trays (Alphamach, DS1922L) (**Fig. S2**). Average temperatures were 0.21 ± 0.28°C (ambient, 1 dpf to 142 dpf) and 4.10 ± 0.48°C (heated, 19 dpf to 100 dpf). A sensor malfunction temporarily stopped heating in the heated reservoir tank, dropping the temperature of the heated incubator trays toward ambient water temperatures. This resulted in the heated incubator reaching 1°C at 83 dpf. This incubator was ramped back up to 4°C by 87 dpf.

Embryos were disinfected biweekly with an immersion in 400 ppm glutaraldehyde water bath in filtered, UV-sterilized seawater for 10 minutes to control potential microbial growth. Dead embryos were removed prior to disinfection and mortality was evaluated by changes in wet weight or by counting individual dead embryos, depending on volume of dead embryos. Every three days, five embryos were randomly sampled and photographed under a dissecting microscope, and their length was measured using an open-source image analysis software platform (ImageJ (103)) with scale bars calibrated to the scope camera.

### RNA extraction and sequencing

To account for different developmental rates at 0°C and 4°C, samples were collected at fixed ages of 30, 60, 90, and 120 days post-fertilization (dpf) and at key developmental milestones heartbeat (HB), 50% eye pigmentation (EP), eye iridescence (EI), and hatching (H). These stages were selected for their clear visibility under a dissecting scope and relatively even distribution throughout development. 120 embryos per time point (30 per technical replicate per treatment) were randomly collected, with chorions pierced for fixative penetration. Whole embryos were preserved in groups of 30 per tube in RNAlater (#AM7201; Invitrogen, Waltham, MA, USA). Embryos were fixed at 4°C for 24 hours, at which point they were moved to −80°C for long term storage. Genomic DNA and total RNA were extracted from single embryos using the Zymo Quick DNA/RNA Kit (#D7001; Zymo, Irvine, CA, USA). Due to their large size (∼12 mm TL), hatchlings were first digested with 20 mg/mL proteinase K solution during a one hour incubation at RT in lysis buffer to ensure full cell dissociation and rupture. RNA extract quality was assessed using an Agilent Tapestation, and only samples with a RNA Integrity Number (RIN) > 6.4 were further processed. Stranded Illumina sequencing libraries were prepared from six replicates per treatment and time point (48 total libraries) at the University of Oregon’s Genomics and Cell Characterization Core Facility (GC3F) using the NuGen Universal Plus mRNA Kit (#0520-A01, Tecan Life Sciences, Männedorf, Switzerland). Resulting libraries were sequenced with paired-end 150 bp reads on an Illumina NovaSeq 6000, averaging 47.3 ± 6.1 million reads per sample (**Table S6**).

### RNA differential expression and variance analysis

The quality of sequenced libraries was visualized using FastQC 0.11.9, where adapter content was seen in some samples. Therefore, adapter trimming was performed with Cutadapt v1.18 (‘--nextseq-trim=18--minimum-length 20’) to remove universal Illumina adaptors (104). Trimmed sequences were aligned to the *Notothenia rossii* genome assembly (GCA_949606895.1; (43)) using HiSat2 v2.2.1 with relaxed mismatch and gap penalties to accommodate polymorphisms between *N. coriiceps* reads and the *N. rossii* genome sequence (parameters ‘--mp 4,1--rdg 4,2 --rna-strandness FR --dta’) (105, 106). On average, 85.5 ± 1.3% of *N. coriiceps* reads aligned to the *N. rossii* assembly (**Table S6**). Transcript annotation of the *N. rossii* assembly derived from the Ensembl Genebuild annotation system (107). Transcript-level expression was estimated with StringTie v1.3.3 (108) and summarized at the gene level using tximport v1.34.0 (109).

Differential expression analysis was conducted with DESeq2 v1.46.0 and EdgeR v4.4.1 (45, 46). Genes were only considered differentially expressed only if they were statistically significant with both tools. Fold change and p-values are reported from DESeq2, with p-values adjusted for multiple testing (FDR). Non-Poisson noise in read count data was estimated using GAMLSS (ExpVarQuant (110)), and the difference in non-Poisson noise between +4°C and 0°C was calculated for each gene. Gene names not found in *N. rossii* annotations were assigned by identifying orthologs in the genome of the channel bull blenny (*Cottoperca trigloides*, GCA_900634415.1, (111)) using BLAST+ v2.14.1 (112).

### Gene ontology enrichment

Since *N. rossii* and *N. coriiceps* lack Gene Ontology (GO) annotations, we used data from the orthologous genes of other vertebrate species. We used OrthoFinder v2.2.6 (113) to identify one-to-one orthologs between *N. rossii* and *C. trigloides*. Genes without orthologs or with many-to-many matches were excluded. GO data were retrieved from *C. trigloides* and Ensembl BioMart-predicted orthologs in human, mouse, chicken, and zebrafish (accessed March 2023) (114). Ensembl gene IDs were then converted into a final, non-redundant *C. trigloides* ortholog set. Functional pathway enrichment was analyzed using clusterProfiler v4.14.4 (48, 115). Certain gene names had more recognizable common names relevant to other taxa. Multiple designations were included for these genes to capture audiences that recognize these genes by different names, rather than only fish related genes.

### Modeling development windows under future climate scenarios

We predicted embryonic viability based on two assumptions supported by the current data: (1) development must occur at ≤ 4°C to complete without abnormalities or hypoxia, and (2) hatching between April and October would increase starvation risk due to limited primary productivity, which is correlated with larval abundance (**Fig. 6A**) (11, 83, 116, 117). To assess future conditions, we used the Coupled Model Intercomparison Project Phase 6 (CMIP6) median sea surface temperature projections for 2100 from the Copernicus Interactive Climate Atlas (**Fig. 6B,C**) and ERA5 data for current conditions (2000–2022) (**Fig. 6B**,**C**) (3, 14).

For each possible fertilization date in the year, we estimated hatching timing and checked for non-permissive developmental temperatures and hatching outside the food resource window. Development duration was assessed using a linear model constrained by known values: 3,672 hours (153 days) at 0°C and 2,088 hours (87 days) at +4°C. The function interpolates the development rate between these points using a linear relationship, expressed as:

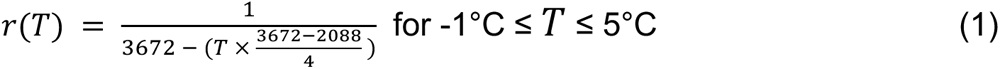

where *r*(*T*) represents the fraction of development completed per hour at a given temperature (*T*). The developmental rate, r(T), was interpolated between −1°C and 5°C to ensure realistic projections. Most current and projected temperatures fall within this range, with modern ERA5 temperatures between −1.62°C and 0.69°C and median of future climate models (SSP5-8.5) between −0.38°C and 5.10°C.

To refine temperature data, we interpolated monthly median temperatures from climate models using a cubic spline method, increasing resolution to an hourly scale. Developmental progress was then computed via numerical integration using the trapezoidal rule with the python function scipy.integrate.cumulative_trapezoid, which approximates the integral as:

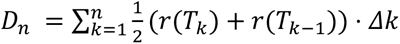

where *D*(*n*) is the cumulative fraction of development completed at time *n*. Hatch timing is calculated by locating the time at which D reaches 1 (i.e., full development). If temperatures exceeded 4°C during development or hatching occurred between April and October, the fertilization date was considered unsuitable.

## Acknowledgements.

We gratefully acknowledge the support of our research by the captain and crew of the *ASRV Laurence M Gould*, the staff of Palmer Station, the personnel of the Office of Polar Programs of the National Science Foundation, and the staff of the Antarctic Support Contractor (Denver, CO). We also offer our heartfelt thanks to Laura Goetz, Sierra Smith, Kathleen Shusdock, Eileen Spann, and Urjeet Khanwalkar whose previous work in 2014 and 2016 field seasons laid the foundation for the success of this study. This work was supported by the NSF grants PLR-1444167 (H.W.D.), PLR-2324998 (J.M.D. and H.W.D.), OPP-1543383 (J.H.P., T.D, and H.W.D.), and OPP-2232891 (J.H.P. and T.D.). This work was completed in part with resources provided by the Research Computing Data Core at the University of Houston.

## Author contributions

M.S., N.R.L.F., T.D., J.H.P., H.W.D., and J.M.D designed research; M.S., N.R.L.F., T.D., J.G., J.H.P, and J.M.D. performed research; M.S., H.W.D., and J.M.D. analyzed data; and M.S. and J.M.D. wrote the paper; M.S., N.R.L.F., T.D., J.G., J.H.P., H.W.D., and J.M.D. provided critical revision of the manuscript.

## Supplemental Figures

**Fig. S1.**
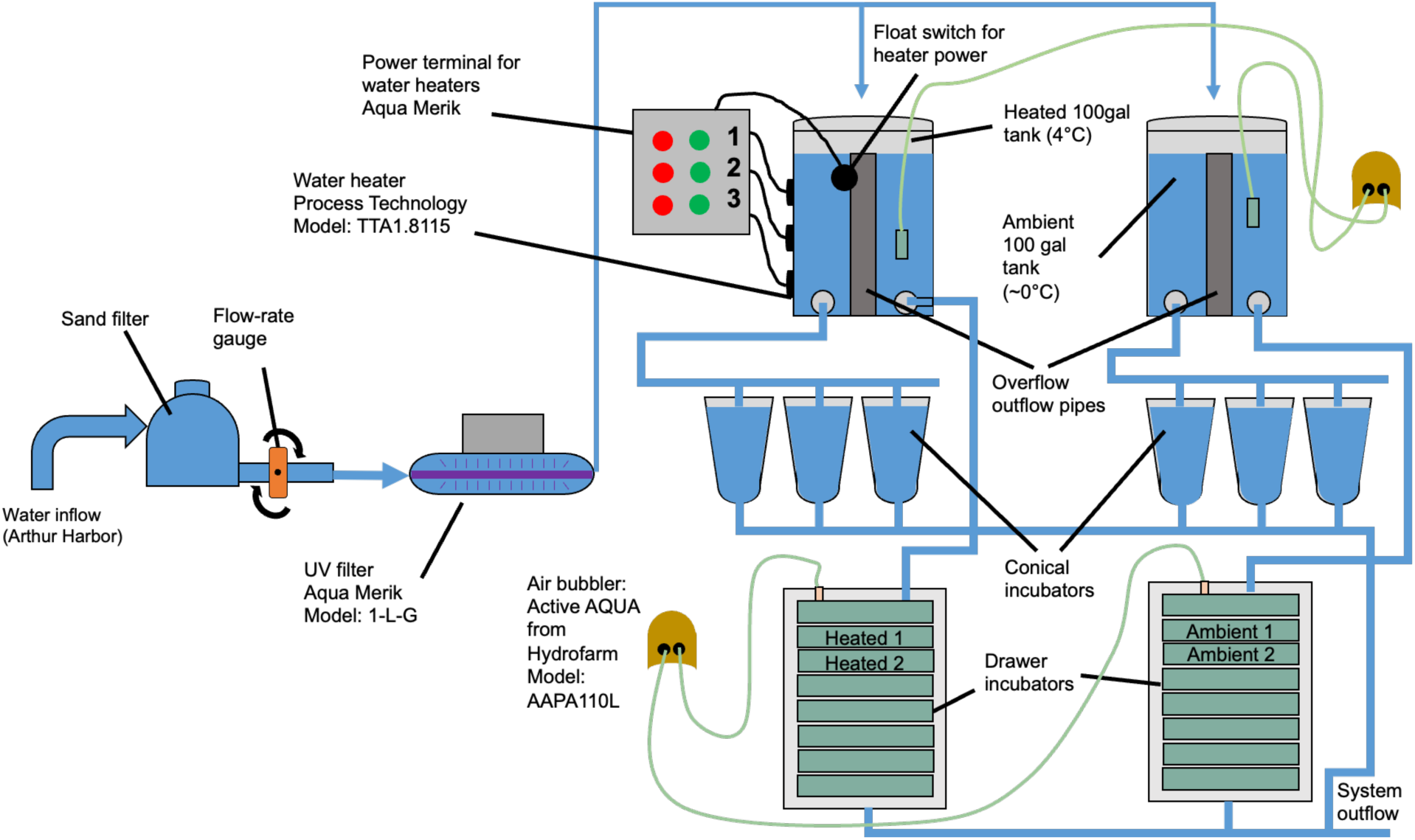
Schematic of vertical incubator system. Seawater entering the Palmer Station Aquarium header tanks was sterilized using a UV light source, and oxygen saturation was maintained with air pumps and spargers. The experimental tank’s seawater temperature was regulated at 4.0 ± 0.2°C using feedback-controlled submersible heaters. The controlled temperature room was maintained at 2°C to keep the control (0°C) seawater near ambient conditions. Seawater flowed to both reservoir tanks and vertical incubators; however, conical jars were not used due to overflow issues but remained connected to the system. Temperature loggers recorded tank temperatures at hourly intervals. Annotations indicate the placement of drawers used for technical replicates.

**Fig. S2.**
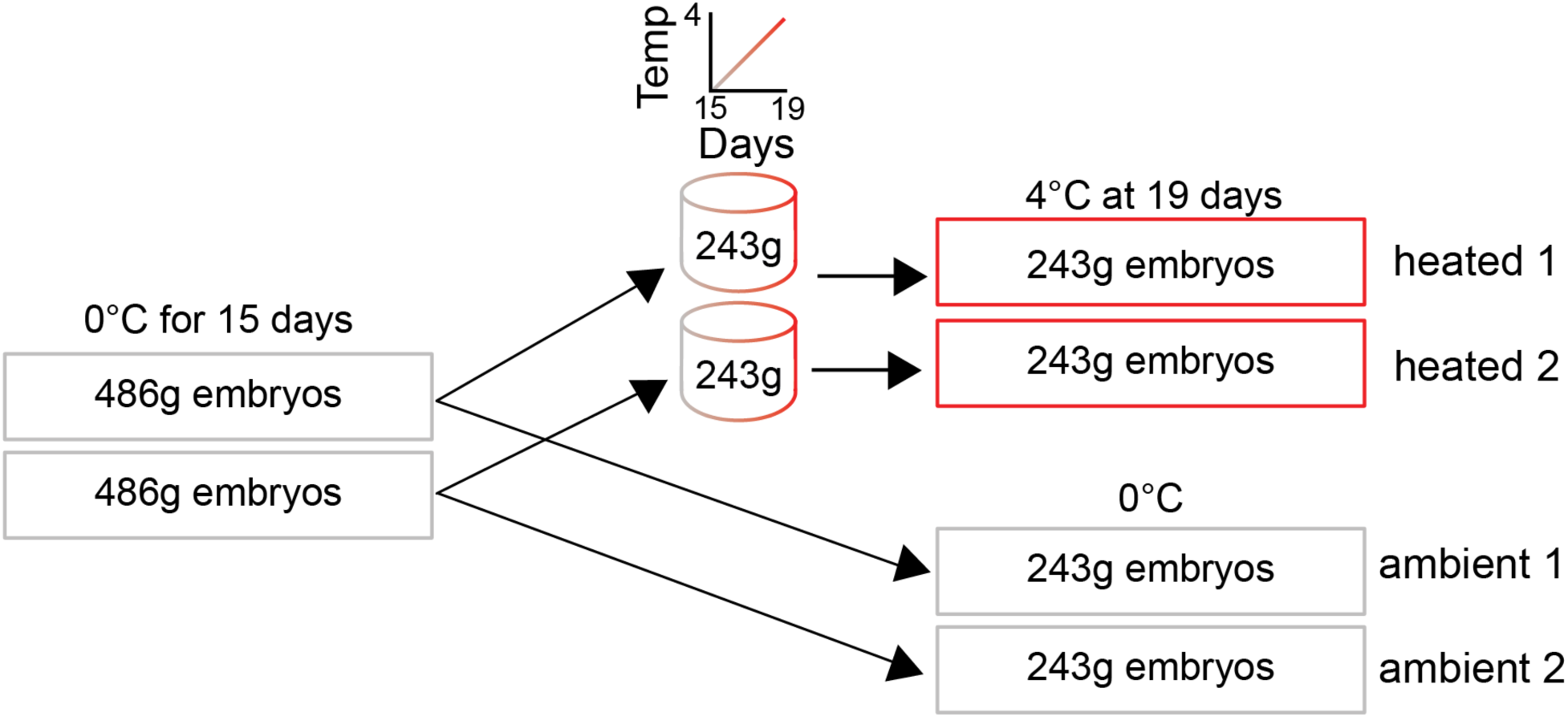
Design of technical replicates. Embryos from four natural, mixed parentage spawns were combined and divided into two incubator drawers (486 g in each). Embryos remained in these incubator drawers until 15 dpf, when half of the embryos were removed from each drawer and moved into large beakers in upright incubators (243g from each replicate tray). The temperature of each upright incubator was raised by one degree per day for four days. At 19 dpf, the embryos from each beaker were placed into two drawers in the heated vertical incubator (Fig S1).

**Fig. S3.**
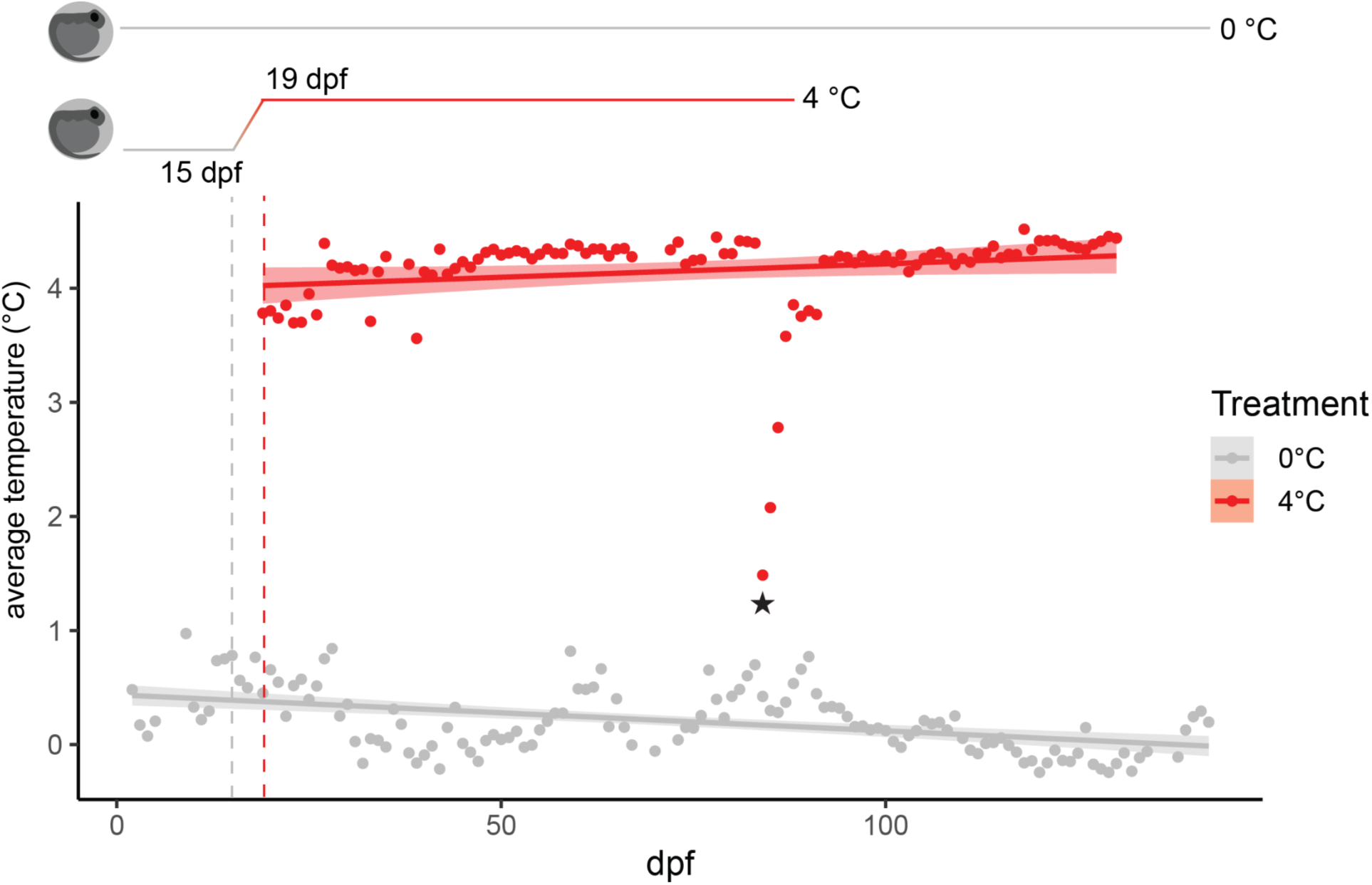
Daily incubation temperatures. Temperatures were recorded hourly using submersed temperature data loggers. Points represent the daily average temperature measured from the bottom drawer of each incubator. The gray dashed line marks 15 dpf, when embryos were removed for a slow temperature ramp to the heated condition (4°C). The red dashed line denotes 19 dpf, when heated embryos were transferred to the heated incubator. Trend lines represent a linear model fitted to each dataset with confidence bands (lm, geom_smooth). The black star at 84 dpf indicates a temperature drop anomaly caused by a malfunction in the flow switch, which temporarily disrupted the heating elements. Temperature was gradually ramped back up by 1°C per day until reaching 4°C.

**Fig. S4.**
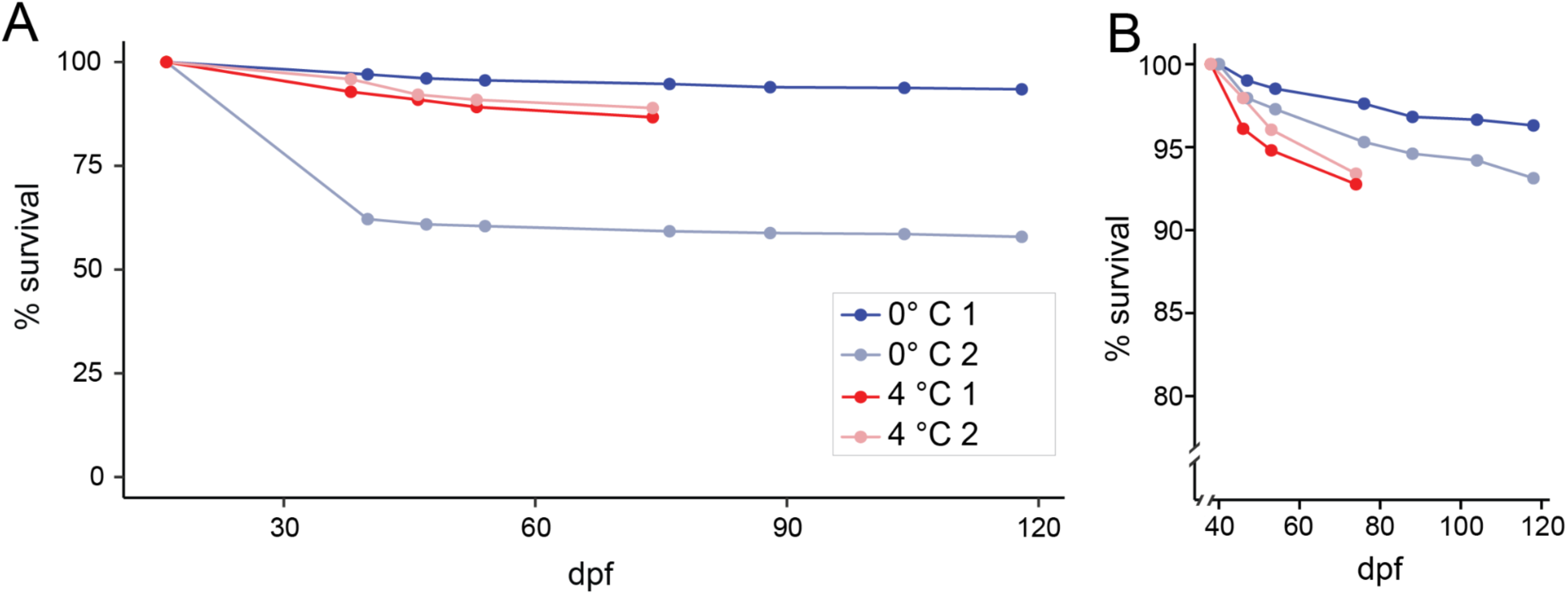
Subtle difference in mortality due to temperature. (A) Percent survival over incubation time, including a mortality event in ambient 2 at 40 dpf. An estimated 1800 embryos died for an unknown reason. However, after this, there was minimal mortality seen during incubation. (B) Percent survival starting from after the mortality event. A slight impact of heat on mortality can be seen, but values were mostly stable, indicating only a subtle difference in mortality due to temperature.

**Fig. S5.**
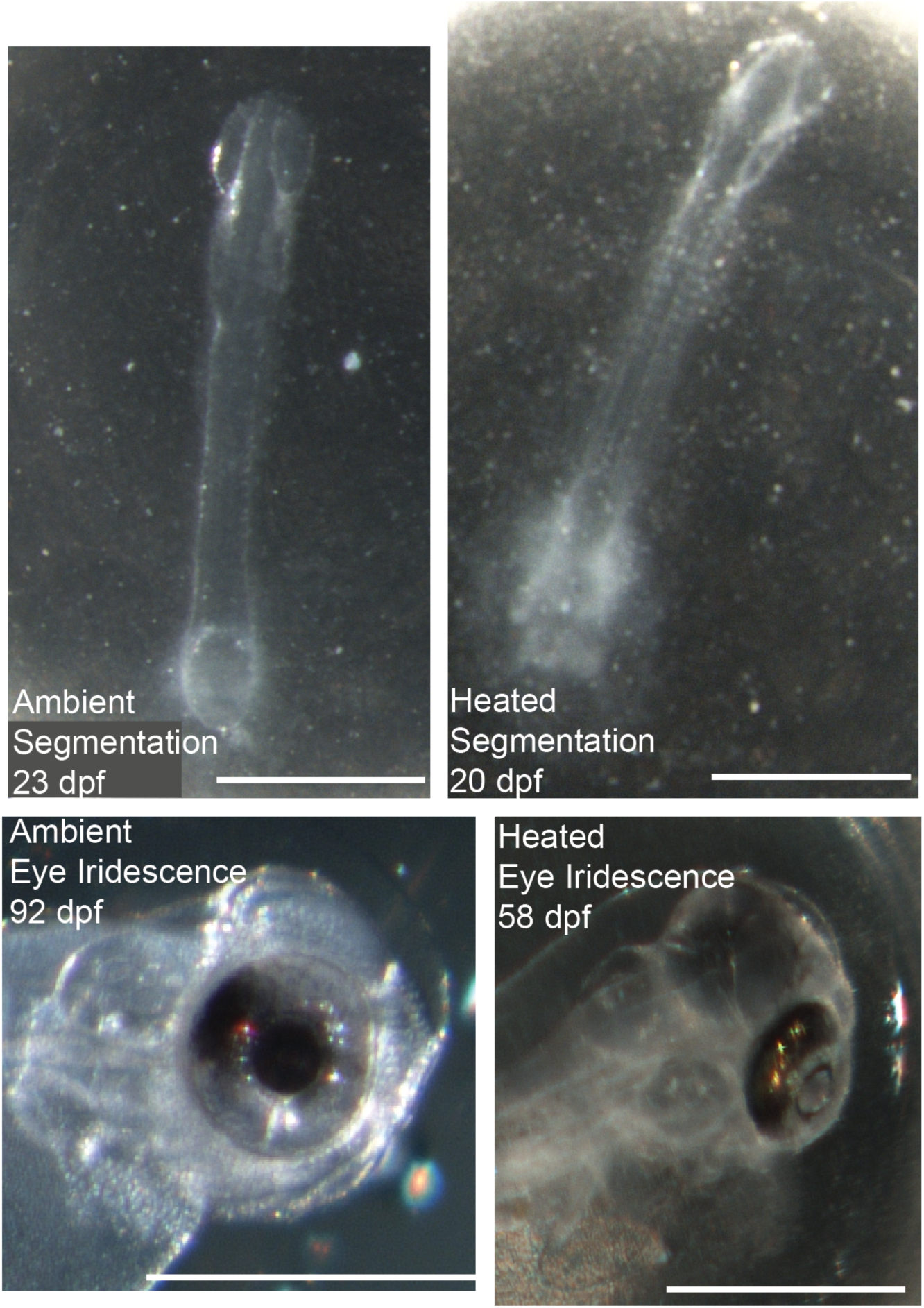
Segmentation and eye iridescence in 0°C and 4°C hatchlings. For eye iridescence, in both cases they were noticed but not immediately photographed. Therefore, the days post fertilization for the photo are not the same as in the text, but several days after. The embryo for ambient eye iridescence was dechorinated before the photograph was taken. All other embryos are pictured within the chorion.

**Fig. S6.**
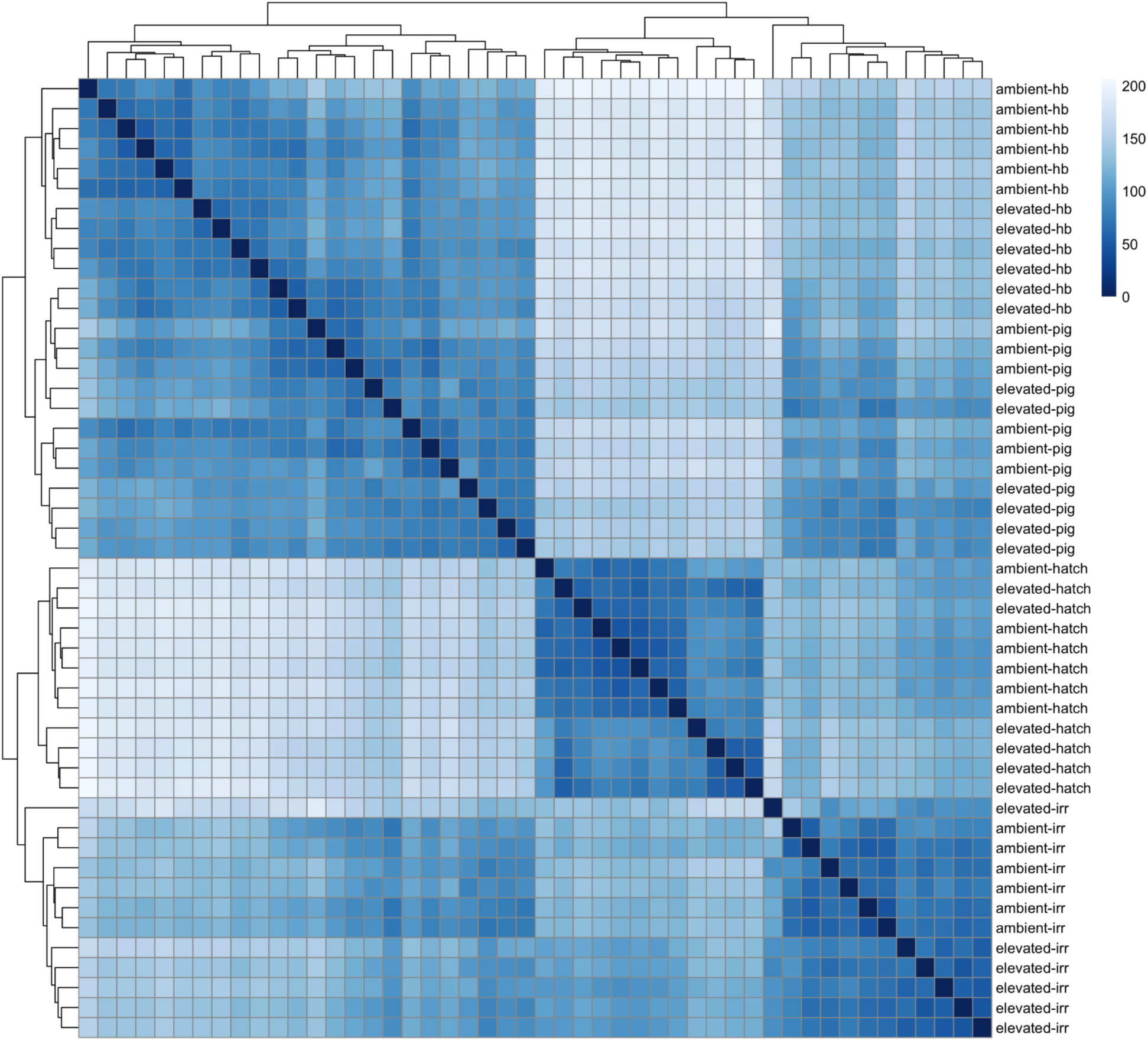
Heatmap of sample distances determined by DESeq2 (v.1.46.0). Samples clustered by developmental stage rather than treatment, indicating consistent gene expression patterns across this morphology-based sampling strategy. Ambient refers to ambient water temperatures (0°C), elevated to 4°C. The analyzed stages are the onset of heartbeat (hb), 50% eye pigmentation (ep), eye iridescence (ir), and hatching (hatch).

## Supplemental Tables

**Table S1.** Timing of developmental stages in 0°C and 4°C treatments.

|  | EXP I (dpf) |  |  |  |
| --- | --- | --- | --- | --- |
|  | Ambient (0C) |  | Heated (4C) |  |
| Heartbeat (HB) | 44 | 7/9/2018 | 34 | 6/29/2018 |
| 50% eye pigmentation (EP) | 65 | 7/30/2018 | 44 | 7/9/2018 |
| Eye iridescence (EI) | 87 | 8/21/2018 | 52 | 7/17/2018 |
| First Hatching (H) | 155 | 10/28/2018 | 87 | 8/21/2018 |

**Table S2.** Measurements and deformity classification of late embryos and hatchlings in 0°C and 4°C treatments. See supporting attachments for spreadsheet.

**Table S3.** Differential expression between treatments of all genes at all stages as reported by DESeq2 v.1.46.0. See supporting attachments for spreadsheet.

**Table S4.** GO enrichment data for all stages. NAs represent terms where no genes were significantly differentially expressed. See supporting attachments for spreadsheet.

**Table S5.**
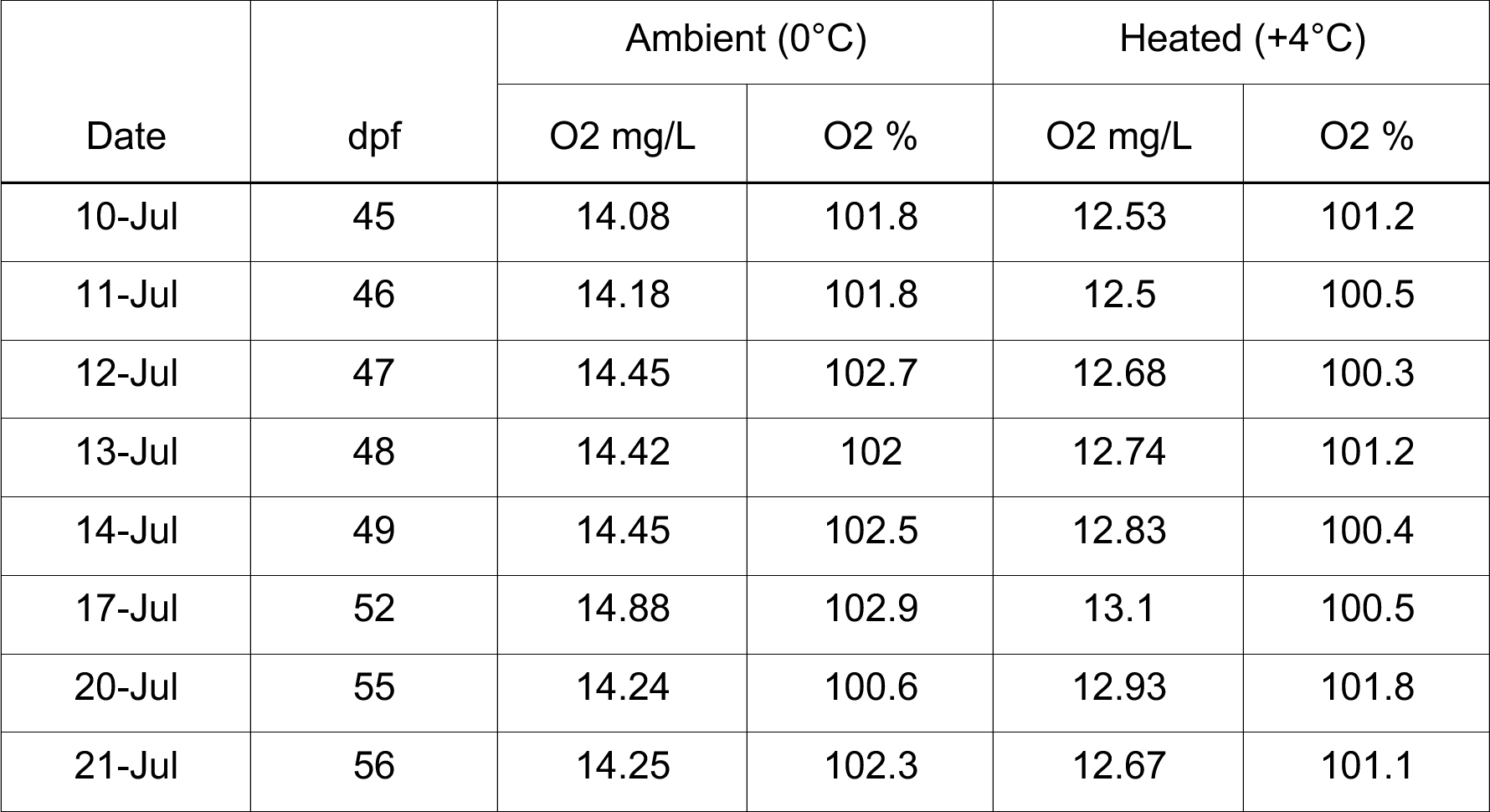
Oxygen saturation in bottom tray of 0°C and 4°C drawer incubators.

**Table S6.** Alignment statistics as reported by Cutadapt v1.18 and HiSat2 v.2.2.1. See supporting attachments for spreadsheet.

